# Parent and offspring genotypes influence gene expression in early life

**DOI:** 10.1101/488684

**Authors:** Daniel J. Newhouse, Margarida Barcelo-Serra, Elaina M. Tuttle, Rusty A. Gonser, Christopher N. Balakrishnan

**Affiliations:** East Carolina University; Indiana State University

**Keywords:** Transcriptome, parental effects, early life stress, nestling, RNAseq, ornithology

## Abstract

Parents can have profound effects on offspring fitness. Little, however, is known about the mechanisms through which parental genetic variation influences offspring physiology in natural systems. White-throated sparrows (*Zonotrichia albicollis*, WTSP) exist in two genetic morphs, tan and white, controlled by a large polymorphic supergene. Morphs mate disassortatively, resulting in two pair types: tan male x white female (TxW) pairs, which provide biparental care and white male x tan female (WxT) pairs, which provide female-biased care. To investigate how parental composition impacts offspring, we performed RNA-seq on whole blood of WTSP nestlings sampled from nests of both pair types. Parental pair type had a large effect on nestling gene expression, with 881 genes differentially expressed (DE) and seven correlated gene co-expression modules. The DE genes and modules up-regulated in WxT nests with female-biased parental care primarily function in metabolism and stress-related pathways resulting from the overrepresentation of proteolysis and stress response genes (e.g. SOD2, NR3C1). These results show that parental genotypes and/or associated behaviors influence nestling physiology, and highlight avenues of further research investigating the ultimate implications for the maintenance of this polymorphism. Nestlings also exhibited morph-specific gene expression, with 92 differentially expressed genes, comprising innate immunity genes and genes encompassed by the supergene. Remarkably, we identified the same regulatory hub genes in these blood-derived expression networks as were previously identified in adult WTSP brains (EPM2A, BPNT1, TAF5L). These hub genes were located within the supergene, highlighting the importance of this gene complex in structuring regulatory networks across diverse tissues.

## Introduction

Parents can have profound impacts on offspring development and fitness. Parental effects can manifest throughout the developmental period, both pre- and post-natally (reviewed in Meaney 2001, Lupien et al. 2009) and can be mediated through parental behaviors, genetics and physiology during early development (Trivers 1972). Parents play a substantial role in establishing the early life environment of offspring. For example in birds, parental decisions on nest placement, incubation behavior, and nest defense could strongly impact developmental conditions of the egg. These parental behaviors will impact exposure to sunlight, humidity, temperature, and other environmental impacts of the eggs, which can influence developmental physiology (e.g. Nord & Nilsson 2011). In addition to parental behaviors, prenatal effects often arise via physiological maternal effects. Developing offspring are susceptible to the maternally created environment (e.g. maternal hormones, immune state, nutrition), which influence offspring physiology (Mousseau & Fox 1998, Jacquin et al. 2012; reviewed in Gluckman et al. 2008, Wolf & Wade 2009, Cottrell & Secki 2009).

The magnitude of parental effects, particularly in altricial species, is likely largest during the postnatal period, when offspring rely entirely on the parents for provisioning and growth (Royle et al. 2012). Provisioning plays a prominent role in offspring development, with the quality and quantity of food items crucial for offspring development (van Oers et al. 2015, Griebel et al. 2019). Similar to the prenatal stage, parental behaviors could also have strong impacts on offspring physiology. In many species, offspring are left alone during parental foraging trips, increasing environmental exposure (Lloyd and Martin 2004) and predation risk (Lima 2009). Parental separation can also increase offspring anxiety (Millstein & Holmes 2007). Siblings must also compete to optimize provisioning, brooding warmth, and preening (Mock & Parker 1997). Thus, this postnatal environment, largely mediated through parental effects, can be a potential source of early life stress (ELS) in offspring, which may result in life-long fitness effects (reviewed in Monaghan 2014).

ELS has broad effects on organisms, including impaired neural development, neuroendocrine signaling, behavior, and physiology (McEwen 2007, Monaghan 2014). For example, ELS is associated with impaired neuroendocrine function and corresponding impaired hypothalamic-pituitary-adrenal (HPA) development, which leads to increase stress response sensitivity later in life (e.g. Heim et al. 2008, Spencer et al. 2009, Crespi et al. 2012, Spencer 2017). ELS can exacerbate behavioral alterations as organisms develop and mature including symptoms of anxiety and depression in the postnatal environment (Noguera et al. 2017) and result in impaired behavior as reproductive adults (e.g. Krause et al. 2009, reviewed in Bolton et al. 2017). While the organismal effects of ELS are well studied, the genetic underpinnings are relatively underexplored. Much of the genetic work in the context of ELS has focused on gene regulatory impacts, particularly in mammalian biomedical models (reviewed in Szyf et al. 2007, Szyf 2009, Silberman et al. 2016, Alyamani & Murgatroyd 2018). In particular, the quality of parental care can have strong impacts on offspring health resulting from epigenetic modifications (Liu et al. 1997, Meaney 2001, Weaver et al. 2004). These gene regulation studies primarily use changes in DNA methylation as an indicator of ELS (Murgatroyd et al. 2009, Kinnally et al. 2011, Lewis & Olive 2014) and recent work has expanded these approaches into non-mammalian organisms (e.g. Rubenstein et al. 2016, Moghadam et al. 2017, Pértille et al. 2017, Gott 2018, Sheldon et al. 2018). DNA methylation studies of ELS investigate changes to the structure of DNA, but are often limited in the functional implications of ELS (i.e. transcription and translation). In general, these modifications are thought to alter transcriptional activity of genes in the modified genomic region (Berger 2007, Lowdon et al. 2016). Indeed, several studies have also taken candidate gene approaches to investigating gene expression in the context of ELS (Marco et al. 2014, Diaz-Real et al. 2017, Anastasiadi et al. 2018, Reshetnikov et al. 2018). However, very few studies assess genome-wide transcription under ELS (Moghadam et al. 2017), particularly in the context of parental effects (but see: Weaver et al. 2006).

In this study, we examined the white-throated sparrow (*Zonotrichia albicollis*, WTSP) to assess the role of parental genotype on offspring gene expression. WTSPs exist in two plumage morphs, tan (T) and white (W), that are found in both sexes and in roughly equal frequencies (Lowther 1961). These morphs are genetically determined by alternative alleles of a supergene, a group of linked genes that are inherited together, show limited recombination, and maintain complex behavioral traits (i.e. WTSP morphs; Schwander et al. 2014, Taylor & Campagna 2016). The WTSP supergene resulted from a complex chromosomal rearrangement comprising multiple inversions (hereafter referred to as “inversion” or “inverted”). This inversion contains ∼1,100 genes on chromosome two, termed ZAL2^m^ (Throneycroft 1975, Thomas et al. 2008, Romanov et al. 2009, Tuttle et al. 2016). W morphs are nearly always heterozygous for the inversion (ZAL2/ZAL2^m^) and T morphs are always homozygous (ZAL2/ZAL2; Thorneycroft 1966, 1975).

This unusual polymorphism in WTSPs influences hormonal profiles and the behavior of both sexes, and thus has the potential to influence pre- and post-natal environments for the offspring of different morphs. W morph males maintain higher levels of testosterone during the pre-laying, incubation, and brooding stages and oestradiol during the laying and brooding stages (Horton et al. 2014). Only oestradiol has been shown to differ between adult female morphs during the breeding season and is higher in W morph females during the pre-laying and laying stages (Horton et al. 2014). These genetic and hormonal differences also translate into striking behavioral differences. W morphs, for example, are highly territorial and sing frequently whereas T morphs are far less territorial and aggressive (Lowther 1962, Kopachena & Falls 1993, Tuttle 2003, Horton & Holberton 2010, Horton et al. 2014). More importantly from the perspective of offspring, males of each morph also differ in paternal investment (Knapton & Falls 1983, Horton et al. 2014). W morph males are promiscuous and provision nestlings very little. T morph males defend their within-pair paternity through mate guarding and are highly paternal. Females tend to provision at intermediate levels, but T morph females may compensate for unassisted care from W morph males and provision more than W morph females (Knapton & Falls 1983). A final wrinkle in this complex mating system is that morphs nearly always mate with the opposite morph (98.5%, Tuttle et al. 2016), resulting in two stable pair types: T male x W female (TxW) and W male x T female (WxT) (Lowther 1961, Tuttle 2003, Tuttle et al. 2016). Because males differ in paternal investment, this results in two distinct parental care strategies. TxW pairs provide biparental care and WxT pairs provide female-biased parental care. In this study we examined gene expression profiles of offspring from both pair-types in order to assess the physiological consequences of variation in parental genotype.

## Methods

### Field based sample collection

All nestling whole blood samples in this study came from a breeding population of WTSPs at the Cranberry Lake Biological Station in northern New York, USA (SUNY-ESF, 44.15°N, 74.78°W) and were collected during the 2015 breeding season. We only utilized samples collected during the first clutch (June 6 - June 14, 2015), as WTSP males may increase paternal investment in replacement broods (Horton et al. 2014). We collected ∼80µL blood in capillary tubes via brachial venipuncture on days 5-7 post-hatch. Approximately 60µL blood was preserved in Longmire’s lysis buffer (Longmire et al. 1992) for genotyping and ∼20µL was immediately placed in RNAlater. Within six hours of collection, samples were placed temporarily into liquid nitrogen, before being shipped overnight on dry ice to -80°C storage until RNA extraction. All animal sampling protocols were approved by the Indiana State University Institutional Animal Care and Use Committee (IACUC 562158-1:ET/RG, 562192-1:ET/RG).

### Molecular sexing & genotyping

Nestling DNA was extracted from erythrocytes using the DNA IQ® magnetic extraction system (Promega Corp, Madison, WI USA). To determine sex and morph, we used PCR to fluorescently label and amplify a region of the chromo-helicase-DNA-binding gene, and a region of the vasoactive intestinal peptide following Griffiths et al. (1998) and Michopolous et al. (2007). The PCR products were run and analyzed on an ABI PRISM™ 310 genetic analyzer.

### RNA extraction, library preparation, & sequencing

We sampled a total of 52 nestlings for RNA extraction, but due to issues with RNA quality after extraction, only 32 were used for sequencing. These samples represent 23 nestlings from eight TxW pairs and nine nestlings from three WxT pairs. Additionally, these data represent 18 females, 14 males, 15 T morph, and 17 W morph individuals.

We removed RNAlater and homogenized whole blood tissue samples with Tri-Reagent (Molecular Research Company). Total RNA was purified with a Qiagen RNeasy mini kit (Valencia, CA, USA), followed by DNase treatment and further purification. We quality assessed RNA with an Agilent Bioanalyzer (RIN > 7) (Wilmington, DE, USA). Both library preparation and sequencing were performed at the University of Illinois Roy J. Carver Biotechnology Center. A library was prepared for each RNA sample using the Illumina HT TruSeq (San Diego, CA, USA) stranded RNA sample prep kit. Libraries were distributed into four pools with equimolar concentrations and quantitated via qPCR. Each of the pools was sequenced on an individual lane of an Illumina HiSeq 2500 using the Illumina TruSeq SBS sequencing kit v4 producing 100-nucleotide single-end reads.

### Creation of masked reference genome

The WTSP reference genome was generated from a male T morph individual (Tuttle et al. 2016). Thus, the reference genome does not contain any sequence data from the ZAL2^m^ inversion. To avoid any potential bias in mapping reads derived from W morph individuals onto a T morph genome, we generated a masked reference genome for this study. To do so, we used previously published whole genome sequences from three W morph adults (Tuttle et al. 2016). Reads were adapter trimmed with *Trim Galore!* v0.3.8 (https://github.com/FelixKrueger/TrimGalore) and aligned to the WTSP reference genome with *bwa mem* v 0.7.10-r789 (Li 2013). We converted and sorted the resulting SAM alignment to BAM format with *samtools view* and *samtools sort*, respectively (*samtools* v1.2, Li et al. 2009). We then merged all genomic scaffolds corresponding to the ZAL2^m^ inversion, as identified in Tuttle et al. (2016), with *samtools merge*. We called SNPs within the inversion using *samtools mpileup* and *bcftools call* v 1.2 (Li et al. 2009, Li 2011). We only kept SNPs that were heterozygous in each of the three individuals with *SnpSift* v 4.3p (Cingolani et al. 2012) and used these SNPs to mask the reference genome with *bedtools maskfasta* v 2.21.0 (Quinlan & Hall 2010).

### Quality control, read mapping, differential expression, & gene ontology

We trimmed Illumina sequencing adapters from each of the 32 libraries with *Trim Galore!* v0.3.8 which uses *Cutadapt* v1.7.1 (Martin 2011). Trimmed reads were then mapped to the masked reference genome with *STAR* v2.5.3a (Dobin et al. 2013). The mapping results were then quantified and assigned gene IDs with *htseq-count* v0.6.0 (Anders et al. 2015) specifying ‘-s reverse’ and ‘-i gene’. We then removed lowly expressed genes by summing the counts for each gene across all 32 samples, dividing by 32 to obtain the study average, and removing genes with an average read count of < 5.

All statistical analyses were performed with R v3.5.0 (R Core Team 2013). We first identified outlier samples based on visual inspection of sample distance in a dendrogram within *WGCNA* (Horvath 2011). Two samples, one T female and one T male representing an entire TxW nest, were identified as outliers and removed from all future analyses (Figure S1). Using the remaining 30 samples, we normalized reads accounting for sequencing depth and assessed differential expression with *DEseq2* (Love et al. 2014). We performed variance stabilizing transformation of reads in *DEseq2* and performed PCA and hierarchical clustering based on Euclidean distance of gene expression profiles with *pcaExplorer* v2.6.0 (Marini & Binder 2016). Differential expression analyses utilized pairwise comparisons between nestling morph and pair type (i.e. parental morphs). We controlled for sex in morph comparisons and sex, morph, and nest ID for pair type comparisons. To include nest ID in the pair type comparison, we followed the “individuals nested within groups” guide in the *DEseq2* manual. We did not include nestling age in analyses, as most samples were 6 days old (n=21), limiting comparisons with nestlings aged Day 5 (n=3) or Day 7 (n=6). Network analysis (see below) did not reveal any effect of age on variables of interest (morph, pair type; data not shown). We also tested for an interaction between nestling morph and pair type utilizing a grouping variable as outlined in the *DEseq2* manual. *DEseq2* determines differential expression with a Wald test followed by Benjamini & Hochberg (1995) FDR correction. Genes were considered differentially expressed (DE) if the FDR corrected p-value was < 0.10. Details for each model run, including the R code used, are in this project’s GitHub repository.

We next tested for gene ontology (GO) enrichment among DE genes with *GOrilla* (Eden et al. 2007, 2009). For each *DEseq2* comparison, we ordered the list of genes based on ascending FDR values, excluding any genes in which *DEseq2* did not assign a FDR value. The WTSP genome is not completely annotated, so any loci without a gene symbol were excluded from GO analyses (n=1,926). *GOrilla* places greater weight on genes located at the top of the list (i.e. DE genes), while accounting for the contribution of each gene in the given comparison. GO categories were considered significantly enriched if the FDR corrected p-value <0.05. *GOrilla* does not support WTSP annotation; so, all analyses were based on homology to human gene symbols.

### Weighted gene co-expression network analysis (WGCNA)

We used the *WGCNA* package in R (Zhang & Horvath 2005, Langfelder & Horvath 2008) to identify modules of co-expressed genes in our dataset. We first exported variance stabilizing transformed (vst) read counts from *DEseq2*, removed genes with an average vst < 5 averaged across all 30 samples, and imported the subsequent list of 8,982 genes into *WGCNA*. To build the co-expression matrix, we chose a soft thresholding power (β) value of 12, at which the network reaches scale-free topology (Figure S2). We generated a signed network with minimum module size of 30 genes and merged highly correlated modules (dissimilarity threshold = 0.20, which corresponds to R^2^ = 0.80). We then correlated the eigengene, which is the first principal component of a module, of these merged modules with external traits (pair type, morph, sex, nest ID). Modules with p < 0.05 were considered significantly correlated with a given trait. For all morph-specific results, we tested for an enrichment of inversion genes with a chi-squared test using a Fisher’s exact test (p < 0.05).

To visualize the interaction of genes within a module, we generated the intramodular connectivity (IM) score for each gene, which represents the interconnection of module genes. We exported all IM scores for modules of interest and imported into *VisAnt* v5.51 (Hu et al. 2013) for visualization. To maximize network clarity, we only plotted the top 300 interactions based on IM scores. Thus, we only visualized the most connected genes. To identify hub genes, we visualized the Degree Distribution (DD) for the network and selected the most connected genes above a natural break in the distribution. This resulted in one to nine hub genes per module.

To understand the biological function of modules correlated with traits of interest, we performed a target vs background GO analysis in *GOrilla*. For each module, we tested the assigned genes for each module against the entire list of 8,982 genes used for the *WGCNA* analysis. GO categories were significant with a FDR corrected p-value < 0.05.

## Results

### Sequencing results

We sequenced each sample to an average depth of 29.4 million reads (range = 16.2-58.5 million reads). The 32 libraries were distributed into four pools in equimolar concentration. One pool contained only four samples, which corresponded to the four samples with lowest RNA concentrations. This pool was sequenced to an average depth of 56.17 million reads per library. The remaining three pools were sequenced to an average depth of 25.62 million reads per library. Samples mapped to our masked genome at an average rate of 91.08% (range = 88.19%-92.87%) (Table S1). A total of 8,982 genes had count values ≥ 5 across all samples, which included 641 located in the W morph inversion. Samples did not segregate by pair type or morph in PCA or hierarchical clustering (Figures S3, S4).

### Differential Expression – Morph

Ninety-two genes were differentially expressed between morphs. Sixty-five of these genes (71%) were located in the inversion, representing a significant enrichment (χ^2^=553.73, df=1, p<0.00001) (Table S2). The inversion represents only 641 out the 8,892 genes (7%) sampled here. Additionally, expression of many of these 92 genes was elevated in W morph nestlings and a number of these genes had well-known functions in innate immunity (e.g. IFIT5, IL20RA, EIF2AK2, RSAD2). There was GO enrichment of four categories, two of which are immunity related: “immune response” (p = 0.019) and “defense response to virus” (p = 0.049) (Table S3).

### Differential Expression – Pair Type

Pair type had the largest effect on gene expression, with 881 genes DE between offspring from the two different pair types (FDR < 0.10, Table S2). Many genes associated with stress responses were elevated in nestlings in WxT nests, including the glucocorticoid receptor (NR3C1), superoxide dismutase (SOD)1 & SOD2, DEP domain-containing mTOR-interacting protein (DEPTOR), and several ubiquitin-mediated proteolysis pathway genes (e.g. UBE2D3, PSMD3, PSMD6). Additionally, several immune system related genes were also elevated in WxT nests, including cytokines (e.g. IL2RA, IL7R), suppressor of cytokine signaling 1 (SOCS1), and five putative major histocompatibility complex (MHC) class I loci. No GO categories were significantly enriched, however.

We next tested for a morph-specific response to pair type. Within WxT nests, 40 genes were DE (p <0.10) between T and W morph nestlings. Twelve of these genes (30%) are located within the inversion, again reflecting an enrichment of inversion genes among those differentially expressed between morph (χ^2^=34.44, df=1, p<0.00001). Only two genes (THSD7B & CFAP44) were DE between morphs within TxW nests, both of which are uniquely DE between morphs in TxW nests. No GO categories were enriched in either comparison.

### WGCNA – Morph

WGCNA revealed 26 modules, five of which were correlated with morph (Table 1, Figure 1). The light cyan module (183 genes, R^2^=0.67, p=5×10^-5^) and ivory module (72 genes, R^2^=-0.66, p=9×10^-5^) contained genes elevated and suppressed, respectively, in W morph nestlings relative to T morph nestlings. These modules are both enriched for genes located within the chromosomal inversion (light cyan module = 70/183 (38%) genes, χ^2^=266.49, df=1, p<0.00001; ivory module = 40/72 (56%), χ^2^=261.60, df=1, p<0.00001) (Figure S5). The hubs of each of these modules are also located in the chromosomal inversion (Table 1, Figure S5). Additionally, the sky blue module (58 genes, R^2^=0.53, p=0.003) and dark red module (102 genes, R^2^=0.47, p=0.009) (Figure S6) contained genes elevated in W morph nestlings and many of these genes overlap with the immune related genes described in the morph DE tests above. The hubs of these networks (e.g. sky blue: EIF2AK2, IFIT5, OASL; dark red: TRAF5) (Table 1) reflect a conserved innate immunity network structure in avian blood (Kernbach et al., in review) (Figure S6).

**Table 1.** WGCNA modules correlated with morph, strength of correlation (R^2^), p-value, hub gene(s) of module, and the degree distribution of hub gene(s).

| **Module** | **$R^2$** | **p-value** | **Hub genes** | **DD of hub gene(s)** |
| --- | --- | --- | --- | --- |
| Dark Red | 0.47 | 0.009 | TRAF5 | 32 |
| Ivory | -0.66 | $9 \times 10^{-5}$ | GOPC, HDAC2, HINT3, TAF5L,<br>TRMT61B, MARC2 | >29 |
| Light Cyan | 0.67 | $5 \times 10^{-5}$ | BPNT1, EPM2A, LOC102066536<br>(GST-like), MAN1A1, MEI4,<br>RNASET2, SLC18B1, TTC32 | >27 |
| Salmon | -0.5 | 0.005 | NSL1 | 39 |
| Sky Blue | 0.53 | 0.003 | DTX3L, EIF2AK2, IFIT5,<br>LOC102064521 (OASL),<br>LOC102065196 (IFI27L2), PARP9,<br>PARP14, RSAD2, ZNFX1 | >22 |

**Figure 1.**
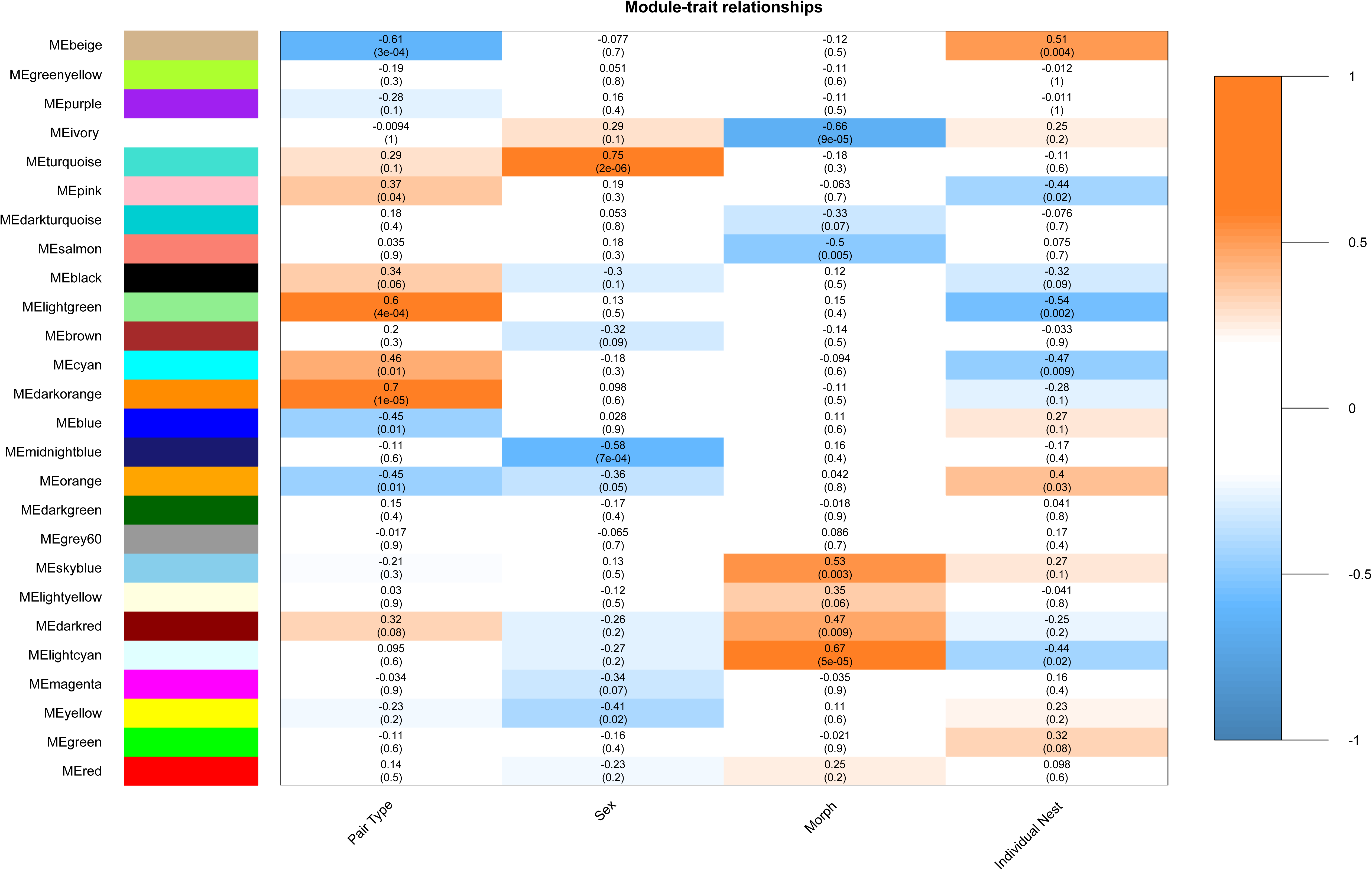
WGCNA module-trait correlation matrix. Each box contains the R^2^ correlation value followed by p-value in parentheses of a given trait with the module. Correlation values range from -1 to 1, with orange colors representing positive correlation and blue colors representing negative correlation.

### WGCNA – Pair Type

We found seven modules correlated with pair type (Table 2, Figure 1). The blue module represented genes that are elevated in nestlings from WxT nests (1,142 genes, R^2^ = -0.45, p=0.01). This module contained both the largest number of genes and correspondingly strongest functional enrichment. Many of these GO enrichments were related to protein function, resulting from the presence of ribosomal genes. Interestingly, several GO categories for metabolism, catabolism, and proteolysis were also enriched, driven by genes encoding ubiquitin-conjugating enzymes and proteasome subunits (e.g. “proteasomal protein catabolic process”, p=2.34×10^-4^; “proteasome-mediated ubiquitin-dependent protein catabolic process”, p=5.32×10^-4^) (Table S4). Many of these (e.g. PSMF1, PSMD3, PSMD6, UBE2D2, UBE2D3, UBE3C) were also DE between offspring of the two pair types (Figure 2). Lastly, the blue module contains one hub gene, NDUFB3 (DD=42) (Figure 2), which is involved in the mitochondrial electron transport chain.

**Table 2.**
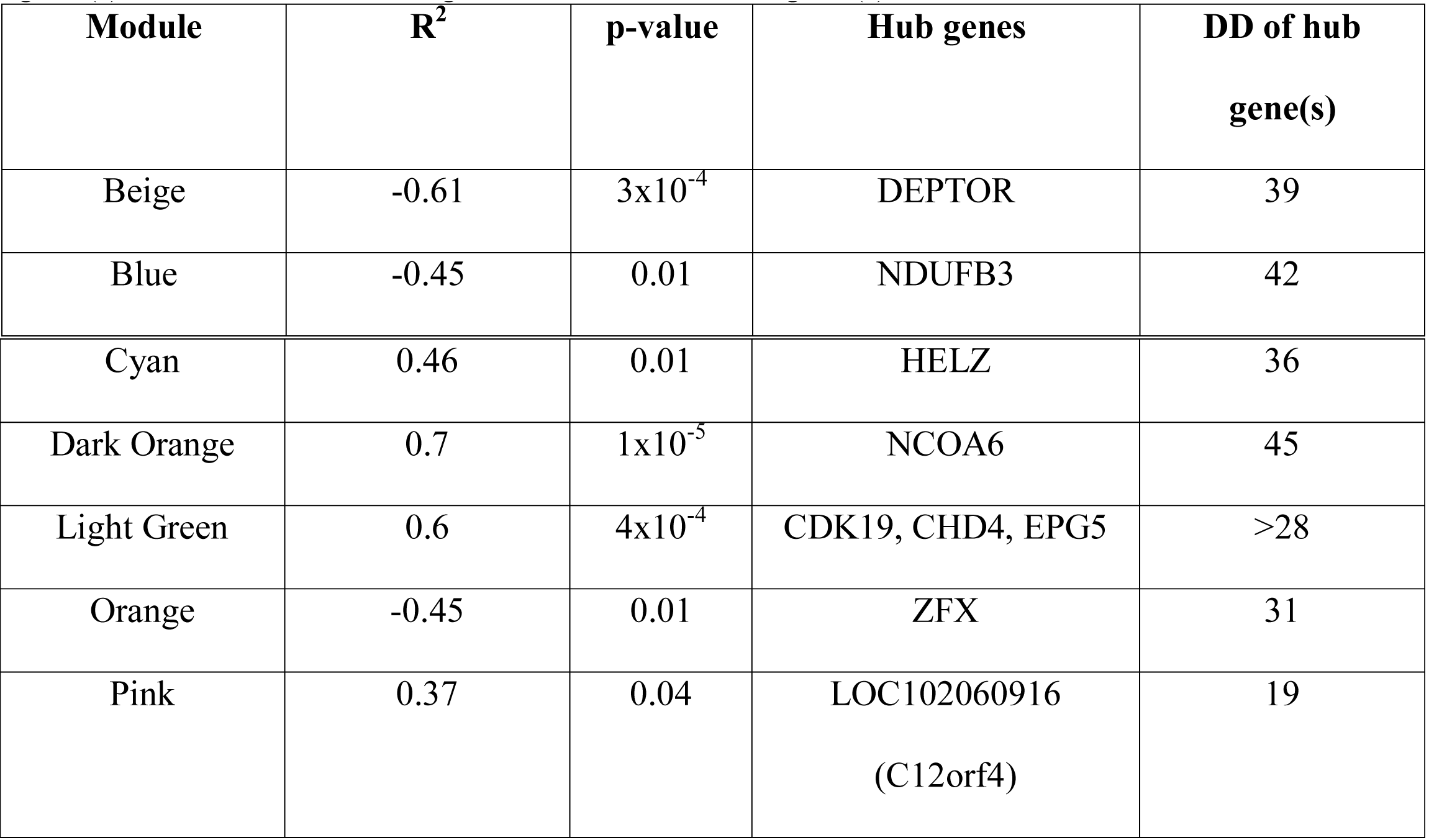
WGCNA modules correlated with pair type, strength of correlation (R^2^), p-value, hub gene(s) of module, and the degree distribution of hub gene(s).

**Figure 2.**
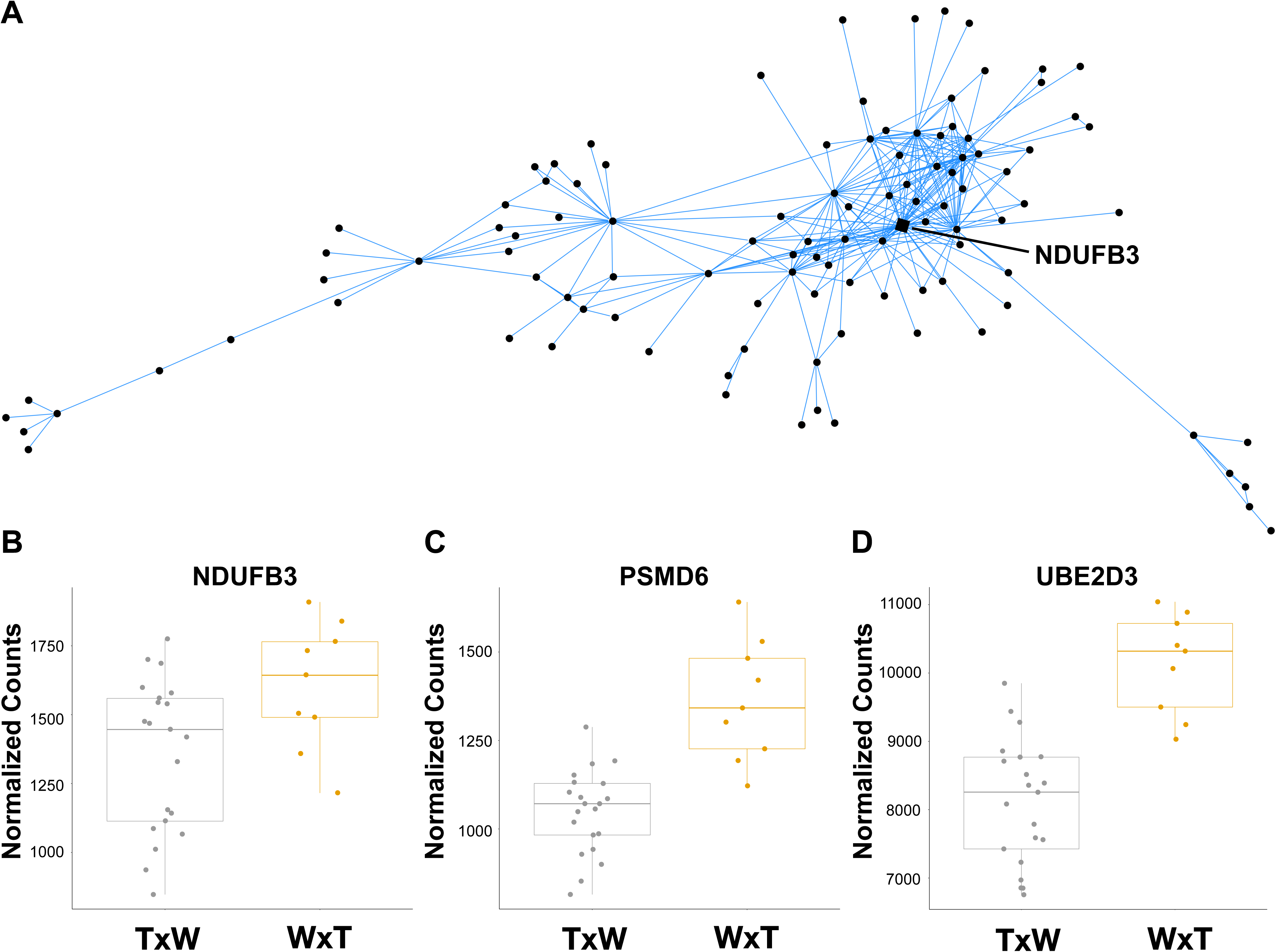
(A) Network of blue module, highlighting hub gene NDUFB3, along with normalized expression plots of (B) NDUFB3, as well as ubiquitin-mediated proteolysis-related genes (C) PSMD6 and (D) UBE2D3. TxW represents samples from nests sired by a T male and a W female. WxT represents samples from nests sired by a W male and a T female. Each circle represents a gene and diamonds represent hub genes described in Table 2.

The beige and light green modules represented candidate stress response networks. These modules showed contrasting expression patterns in nestlings from WxT nests (Figures 4 & 5). Although not significantly enriched for any GO categories, the beige module comprised 335 genes that were upregulated in WxT nests relative to TxW nests (R^2^=-0.61, p=3×10^-4^). DEPTOR, which functions as an inhibitor of the mTOR pathway in response to stress (e.g. Desantis et al. 2015), was the single hub in the beige module (DD=39, Figure 3). The beige module also contained NR3C1, which is activated in response to increased glucocorticoid secretion. Lastly, the light green module (116 genes, R^2^=0.60, p=4×10^-4^) contained genes with low expression in TxW nests relative to WxT nests. There were three hub genes (DD > 28), CDK19, CHD4, and EPG5, each with previously described roles in the stress response (Figure 4).

**Figure 3.**
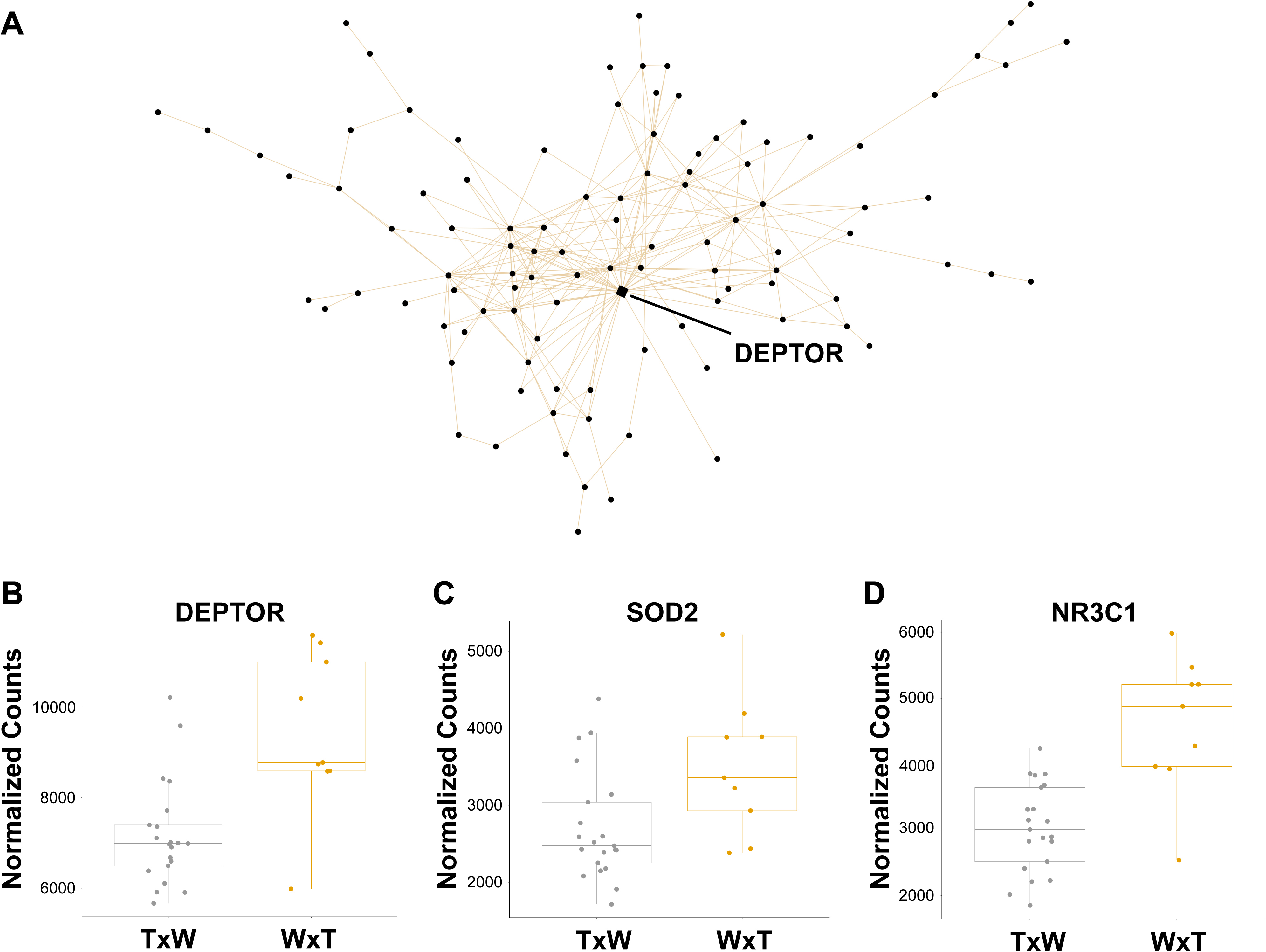
(A) Network of beige module, highlighting hub gene DEPTOR, along with normalized expression plots of hub gene (B) DEPTOR, as well as stress responsive genes (C) SOD2 and (D) NR3C1. TxW represents samples from nests sired by a T male and a W female. WxT represents samples from nests sired by a W male and a T female.

**Figure 4.**
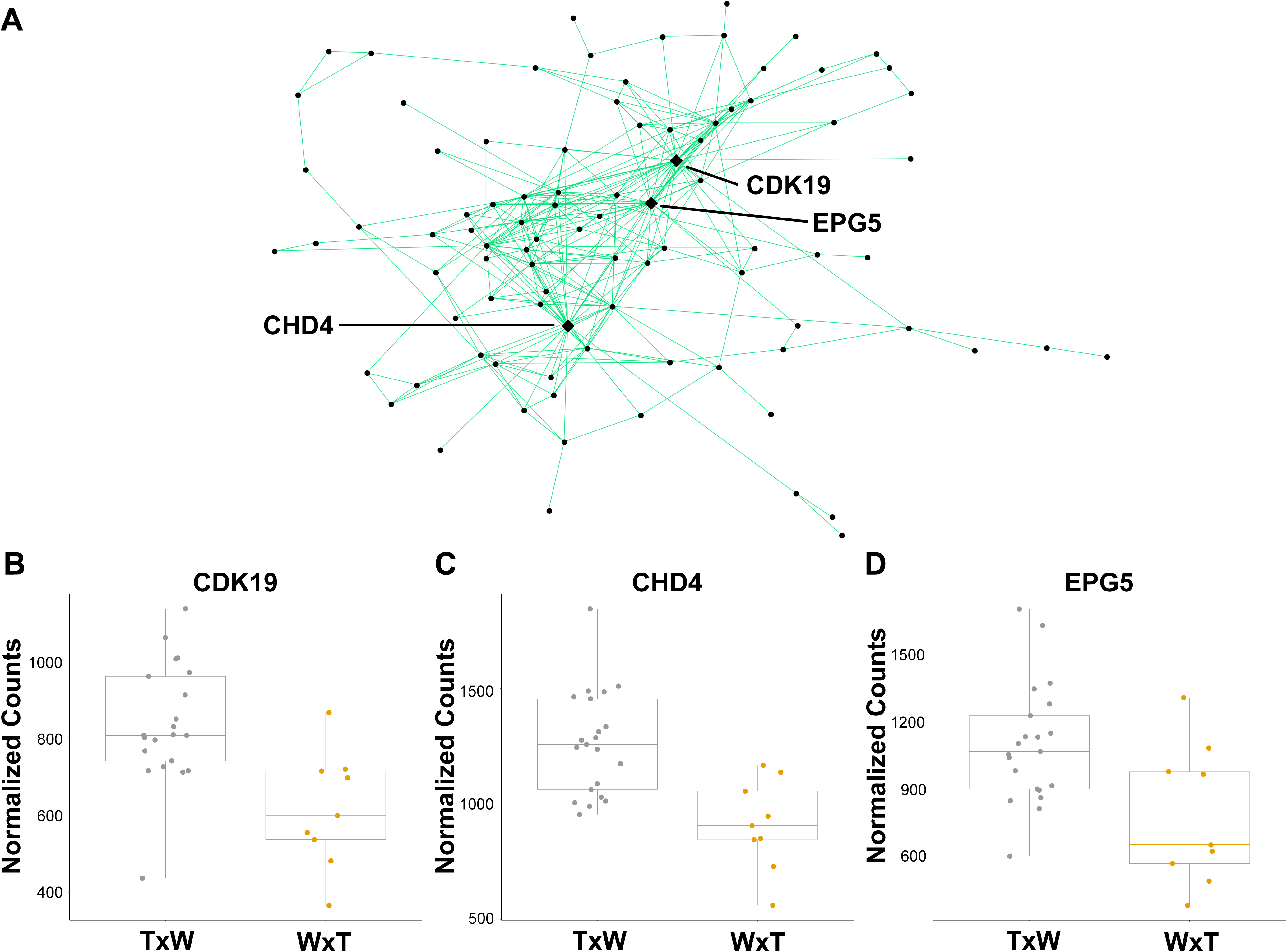
(A) Network of light green module and normalized expression plots of hub genes (B) CDK19, (C) CHD4, and (D) EPG5. TxW represents samples from nests sired by a T male and a W female. WxT represent samples from nests sired by a W male and a T female. Each circle represents a gene and diamonds represent hub genes described in Table 2.

For each pair type module, the correlation was stronger for the overall effect of pair type than any individual nest, indicating that one nest did not drive the correlation. This trend was reflected in gene expression plots of hub genes and candidate genes described above (Figure S7). We did not observe modules correlated with pair type that were also correlated with nestling morph or sex, suggesting there is no morph or sex-specific response to a given pair type at the network level.

## Discussion

By assessing genome-wide transcription in nestlings raised by different WTSP pair types we have identified distinct transcriptomic signatures that suggest WxT pairs induce a stress response in developing nestlings relative to TxW pairs. This is reflected both by differential expression of several genes involved in protein degradation as well as networks of co-expressed genes with stress response hubs. Additionally, we identified morph-specific gene expression driven by innate immunity genes and genes located in the chromosome 2 inversion. As adults, the genes within the inversion strongly influence the WTSP neural transcriptome (Balakrishnan et al. 2014, Zinzow-Kramer et al. 2015). Our results here suggest that as nestlings, parental genotypes and associated behaviors, rather than nestling genotype, have the strongest influence on the nestling transcriptome.

### Gene expression differences resulting from pair type

We find 881 genes DE between nestlings raised under the two pair types. Many of these genes function in the proteasome or ubiquitin-mediated proteolysis. Cells naturally use the proteasome for degradation of proteins targeted by the ubiquitination process, but genes involved in proteasome formation (e.g. PSMD6, PSMD11) and ubiquitination (e.g. UBE2B) are up-regulated in cells experiencing mild oxidative stress (Aiken et al. 2011, Shang & Taylor 2011, Livneh et al. 2016) or organisms experiencing abiotic stress (Dhanasiri et al. 2013, Tomalty et al. 2015). Thus, increased expression of these genes in nestlings from WxT nests suggests they are responding to oxidative stress. As a result, there may be a cost to having a W morph father and T morph mother at the nestling stage.

To complement our differential expression approach, we also constructed co-expression networks with *WGCNA. WGCNA* identifies modules of co-regulated genes blind to the experimental design. These modules are then correlated with external traits, offering a systems-level view into how conditions impact transcriptional networks. Within these networks, we can then perform GO analyses as described above and identify network hubs, which are the most highly connected genes within that network. Using this approach, we identified 26 modules of co-regulated genes in this dataset (Figure 1), seven of which were significantly correlated with parental pair type. The blue module contains genes that are elevated in nestlings in WxT nests. The blue module hub gene was NDUFB3 (Module Membership [MM]=0.938, DD=42) (Figure 2), which encodes a subunit of the mitochondrial membrane respiratory chain. Interestingly, many of the same proteolysis-related genes highlighted in the differential expression results are also present in this module, resulting in the enrichment of several metabolism and stress-related GO categories (Table S4).

Two modules, light green and beige, contained stress responsive hub genes. The light green module contains genes that are suppressed in nestlings in WxT nests, with three hub genes: CDK19, CHD4, and EPG5 (Figure 4). The absence of EPG5 expression (via knockout) and reduction in CHD4 expression (via knockdown) has been associated with increased DNA damage (Zhao et al. 2013, Larsen et al. 2010). Similarly, down-regulation of CDK19 following knockdown is associated with an increased stress response (Audetat et al. 2017). Suppression of these genes in these nestlings could be indicative of increased cellular damage. The beige module contains genes whose expression is elevated in nestlings from WxT nests and contains one hub gene, DEPTOR, which is an inhibitor of mTOR signaling (Figure 3). The exact role of DEPTOR remains unclear, but up-regulation likely inhibits the mTORC1 pathway to reduce endoplasmic reticulum stress, promote cell survival, and avoid apoptosis (Peterson et al. 2009, Desantis et al. 2015, Catena et al. 2016).

Increased expression of genes in the beige module in these nestlings and the high connectivity of DEPTOR to other co-expressed genes provide further support for a transcriptional stress response within WxT nests. The beige module also contains two well-studied stress responsive genes, superoxide dismutase 2 (SOD2) and the glucocorticoid receptor (NR3C1). SOD2 mitigates the effects of exposure to reactive oxygen species by scavenging free radicals (Zelko et al. 2002). NR3C1 binds glucocorticoids and has primarily been studied in the context of ELS and methylation of an upstream promoter. NRC3C1 methylation is often associated with down-regulation of NR3C1 (e.g. McGowan et al. 2009) and impairment of the HPA axis, but up-regulation following methylation has also been observed as part of the stress response (Turner et al. 2006, Bockmühl et al. 2015). Increased expression observed here directly implicates the HPA axis and suggests these nestlings may be activating SOD2 and NR3C1 to cope with elevated levels of reactive oxygen species and corticosterone, respectively (Wang et al. 2018, Finsterwald & Alberini 2014). However, further work is needed to investigate stress physiology, corticosterone levels, and uncover the epigenetic state of NR3C1 in these nestlings and how this may relate to ELS (Banerjee et al. 2011, McCoy et al. 2016, Rubenstein et al. 2016, Quirici et al. 2016, Greggor et al. 2017).

### How does parental genotype influence offspring gene expression?

In a non-experimental study, we have limited power to make inferences about the mechanism by which parental genotype impacted offspring gene expression. Given the well-studied reproductive biology of WTSPs, however, two mechanisms seem especially likely: hormone-mediated maternal effects and/or differences in parental provisioning. In weighing the evidence for these two non-mutually exclusive possibilities, we conclude that the difference in parental provisioning is the most plausible explanation for the observed gene expression differences. As described above, WTSP morphs differ in hormone levels. Only oestradiol, however, has been shown to differ between adult female morphs during the breeding season and is higher in W morph females during the pre-laying and laying stages (Horton et al. 2014). No baseline differences in any other hormone measured to date (corticosterone, testosterone, DHEA, DHT) have been described during the breeding season (Spinney et al. 2006, Swett & Breuner 2009, Horton & Holberton 2010, Horton et al. 2014). Taken together this suggests that hormone deposition into eggs may not differ dramatically between the morphs. By contrast, there is strong evidence of differences in provisioning among morph types (Knapton & Falls 1983, Kopachna & Falls 1993, Horton & Holberton 2010, Horton et al. 2014). Reduced provisioning by W morph males appears to be stable across populations resulting in female-biased parental care in WxT nests (Knapton & Falls 1983, Horton et al. 2014). Therefore, parental care variation is a likely source of behaviorally mediated maternal or paternal effects (see Crean & Bonduriansky 2014) that would explain the strong signature of stress exposure in the expression data.

Previous work revealed no difference in clutch size between pair types (Knapton et al. 1984, Formica et al. 2004) and no effect of pair type on nestling mass (Knapton et al. 1984, Tuttle et al. 2017). Also, nestlings did not differ in mass at time of sampling between the TxW and WxT nests used in this study (Smith et al. in review). Increased provisioning by females to compensate for reduced care by males could explain this observation, and this has been observed previously in a separate WTSP population (Knapton & Falls 1983). In this scenario reduced brooding and increased maternal separation could also negatively impact nestling physiology and act as a source of ELS (reviewed in Ledón-Rettig et al. 2013). Somewhat surprisingly, given the gene expression findings described here, a recent study in our study population did not detect differences in reactive oxygen metabolites in plasma of offspring of the two different pair types (Grunst et al. 2019). ROM, however, only provides a limited overview of the stress response and the RNA-seq response we observed could even mitigate long-term consequences of ELS. The results here further highlight the utility of blood RNA-seq as a highly sensitive measure of environmental exposures (Louder et al. 2018).

Our study was carried out in the field as part of a long-term study and is limited by the fact that we did not perform a cross-fostering experiment. We aimed to mitigate potential environmental confounds by restricting sampling of nestlings to a short time period of nine days. Certainly the environment may influence gene expression in our samples, but consistent changes among the samples in the two pair types suggest the role of parents is a significant driver of nestling gene expression, rather than temporal or spatial environmental variation.

### Morph-specific gene expression

We were also interested in morph-specific gene expression and how nestling morph may respond to differences in parental pair type. WTSPs have been studied extensively as adults, but very rarely in other life stages. W morph males and T morph females exhibit earlier reproductive and actuarial senescence, potentially resulting from the high energy expenditure lifestyle of W morph males and biased parental care given by T morph females (Grunst et al. 2018a, Grunst et al. 2018b). There also appears to be seasonal variation in fitness between the morphs as adults. Following cold, wet winters, W morph males exhibit lower recruitment in the breeding grounds, leading to an overproduction of W morph male nestlings, potentially to stabilize morph frequencies in the population (Tuttle et al. 2017). Thus, morph specific differences may arise in early life. We found 92 genes DE between morphs, including many innate immune-related genes and genes located within the inversion (65/92 genes, Table S2). *WGCNA* revealed five modules correlated with morph (Figure 1). These included two innate immunity-related modules with increased expression in W morphs (Dark Red & Sky Blue) and two modules enriched with genes located in the inversion (Ivory = 40/72, Light Cyan = 70/183) (Figures S5, S6). The sky blue module contains nine hub genes and the dark red module contains one hub gene, both of which include well-studied anti-viral genes (e.g. sky blue: OASL, RSAD2; dark red: TRAF5). These genes also form a co-expression module in avian blood following West Nile virus infection (Kernbach et al., in review). Adult WTSP morphs differ in their ability to clear infection (Boyd et al. 2018), so the immune activation here may be indicative of an increased parasite load in W morph nestlings, although further investigation is required. The light cyan module contains genes elevated in W morph nestlings and contains eight hub genes, each located in the inversion (Table 1). Three of these, EPM2A, BPNT1, and TAF5L, were also identified as hub genes in brain tissues of adult W morph males (Zinzow-Kramer et al. 2015). These nestlings thus exhibit expression differences in inversion genes prior to any phenotypic or behavioral differences, revealing the importance of the inversion in maintaining morph phenotypes throughout life. Additionally, the conservation of network hub genes in a different tissue and life stage highlights avenues for further investigation into WTSP gene regulation.

Despite broad gene expression differences between the morphs, within pair types morph-specific expression was limited. In part due to small sample size, nestlings in TxW nests only have two genes DE between morphs. There is a larger effect of morph within WxT nests, where the number of DE genes increased to 40. These genes encompassed a wide range of gene functions without any obvious stress-related candidate genes. Of these 40 genes, 34 are uniquely DE within WxT nests and do not overlap with the overall list of 92 genes DE between morphs using all samples. Interestingly, glucocorticoid-induced transcript 1 (GLCCI1) is elevated in W morph nestlings in WxT nests. The function of GLCCI1 remains unclear (Kim et al. 2016), but expression differences between morphs observed here implicates the role of glucocorticoids in response to pair type. This suggests that nestling morphs may respond differently to the parental pair type though larger sample sizes will be needed to explore this further.

## Conclusions

Using the WTSP, a system with alternative parental care strategies, we show that nestlings in WxT nests (female-biased parental care) have increased expression of stress-related genes, and parental genotypes may act as a source of ELS in the species. Nestling morph also influences transcription, but parental pair type appears to have the greatest effect on their transcriptome. Combined, this supports the parental effects hypothesis (Wade 1998, Schrader et al. 2018), where offspring phenotypes are primarily a result of the nest environment and care received, rather than from offspring genotypes (i.e. T vs. W). Nearly 54% of observed pairs have been WxT (Tuttle et al. 2016). Thus, roughly half of the nestlings in every population will experience female-biased parental care. Our results suggest that these differences in parental pair type have at least short-term consequences on offspring physiology. While we have identified impacts at the level of transcription, an integrative approach assessing nestling WTSP physiology and performing cross-fostering experiments will further elucidate the consequences of variation in parental pair type. Importantly, it remains unclear whether female-biased parental care or differences in maternal effects translate into long-term fitness consequences for offspring. There appears to be a cost associated with parental genotype, as less cooperative reproductive strategy (WxT pairs) accelerates senescence (Grunst et al. 2018a, Grunst et al. 2018b). We show here that this cost is also translated into nestlings within WxT nests via increased stress-related gene expression. This work sets the stage to further explore morph-specific fitness consequences in nestlings experiencing alternative parental care strategies.

## Supporting information

Table S2: DEseq2 Results

Table S4: WGCNA Results

## Acknowledgements

We acknowledge Lindsay Forrette, Andrea Grunst, and Melissa Grunst for assistance in the field, Sarah Ford for assistance with molecular work, Rachel Wright for WGCNA code, and Cranberry Lake Biological Station. Funding provided by East Carolina University, Indiana State University, The National Science Foundation (grant no. DUE-0934648) and the National Institutes of Health (grant no. 1R01Gm084229 to E.M.T and R.A.G.) and a Sigma Xi Grants in Aid of Research award to DJN. Birds were banded with color bands and a Fish and Wildlife band (Master Banding Permit 22296 to EMT and permit 24105 to RAG). Dr. Alvaro Hernandez and Chris Wright provided guidance and oversight on sequencing carried out at the University of Illinois. All methods were conducted in accordance with legal and ethical standards and were approved by Indiana State University’s Institutional Animal Care and Use Committee (protocols 562158-1:ET/RG and 562192-1:ET/RG).

## Data Accessibility

The 32 RNAseq libraries used in this study will be submitted to the NCBI Sequence Read Archive (SRA). All files needed to produce these results, including code and counts files, will be uploaded to this project’s GitHub page: https://github.com/danielnewhouse/wtsp

## Author Contributions

DJN designed and performed research, analyzed the data, and wrote the paper. MBS performed research, contributed samples, and reviewed drafts of the paper. EMT designed and performed research and contributed samples. RAG designed and performed research, contributed samples, and reviewed drafts of the paper. CNB designed and performed research, contributed reagents, and reviewed drafts of the paper.

