## Supplemental Tables and Figures for "Parent and offspring genotypes influence gene expression in early life"

| Sample | Mapping rate |
|---|---|
| 25898 | 88.19 |
| 37919 | 88.73 |
| 25880 | 88.77 |
| 25878 | 89.09 |
| 37923 | 89.14 |
| 25889 | 89.73 |
| 25876 | 90.24 |
| 37902 | 90.47 |
| 37917 | 90.62 |
| 25875 | 90.83 |
| 37906 | 90.94 |
| 37915 | 90.94 |
| 37927 | 90.94 |
| 37916 | 90.98 |
| 37918 | 90.98 |
| 37913 | 91.24 |
| 25881 | 91.48 |
| 37930 | 91.6 |
| 37925 | 91.64 |
| 37901 | 91.71 |
| 25877 | 91.79 |
| 25896 | 91.89 |
| 37929 | 91.91 |
| 25883 | 91.93 |
| 25888 | 91.93 |
| 37926 | 91.94 |
| 25900 | 91.98 |
| 37933 | 92.21 |
| 25882 | 92.63 |
| 37922 | 92.67 |
| 25890 | 92.74 |
| 25887 | 92.87 |
| Average | 91.0859375 |

**Table S1.** Mapping rates to the masked reference genome for each sample used in this study.

**Table S2.** Differential expression results for morph and pair type contrasts. The results are provided in an Excel spreadsheet available on the journal website.

| GO term | Description | P-value | FDR q-value | Enrichment (N, B, n, b) |
| --- | --- | --- | --- | --- |
| **GO:0006955** | immune response | 1.55E-06 | 1.89E-02 | 2.16 (6812,242,625,48) |
| **GO:1903047** | mitotic cell cycle process | 4.82E-06 | 2.95E-02 | 1.77 (6812,342,821,73) |
| **GO:0080134** | regulation of response to stress | 1.54E-05 | 6.27E-02 | 2.77 (6812,560,114,26) |
| **GO:0051607** | defense response to virus | 1.59E-05 | 4.86E-02 | 5.11 (6812,57,304,13) |

**Table S3.** Gene Ontology results for the morph differential expression contrast. Enrichment (N, B, n, b) is defined as follows: N - is the total number of genes, B - is the total number of genes associated with a specific GO term, n - is the number of genes in the top of the user's input list or in the target set when appropriate, b - is the number of genes in the intersection, Enrichment = (b/n) / (B/N).

**Table S4**. WGCNA results with module assignment and module membership score for each gene. Gene Ontology results provided for modules of interest. The results are provided in an Excel spreadsheet available on the journal website.

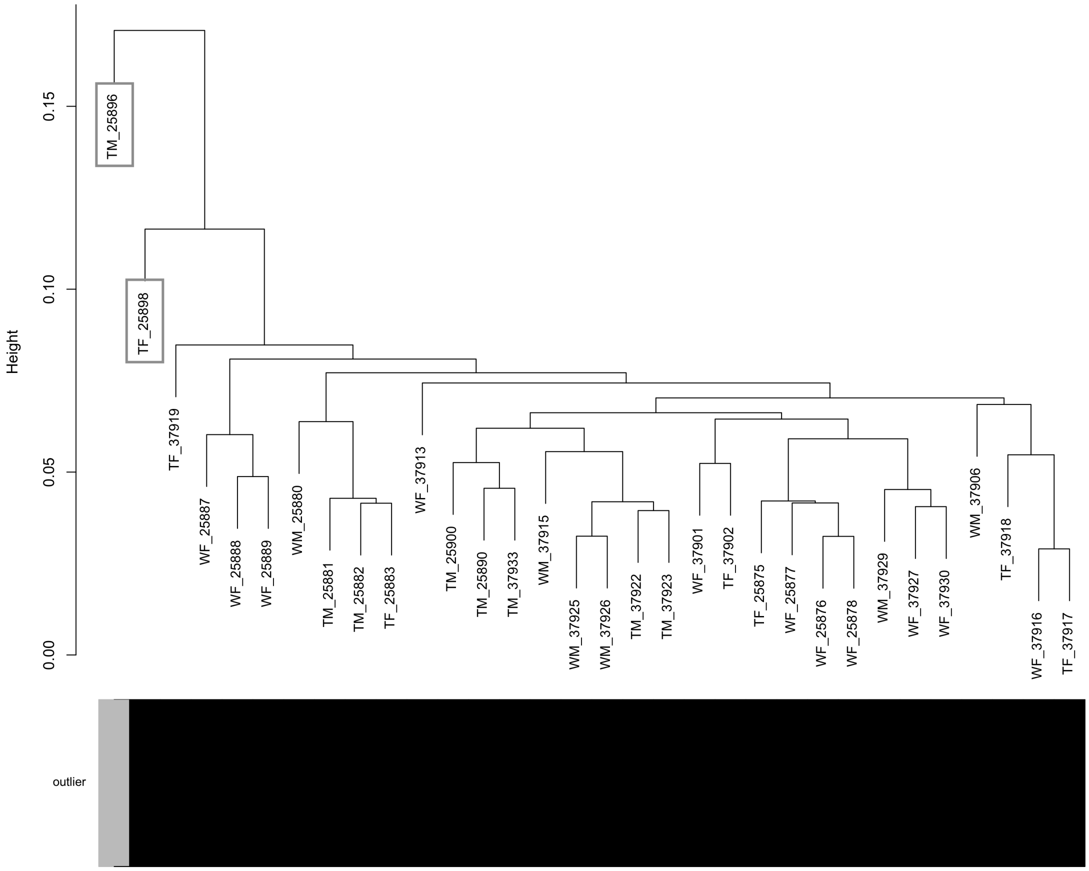


**Figure S1**. Sample dendrogram showing two samples, TM_25896 and TF_25898, identified as outliers and removed from analyses. Samples are named by morph/sex combination followed by sample ID. TM denotes tan male, TF denotes tan female, WM denotes white male, and WF denotes white female.

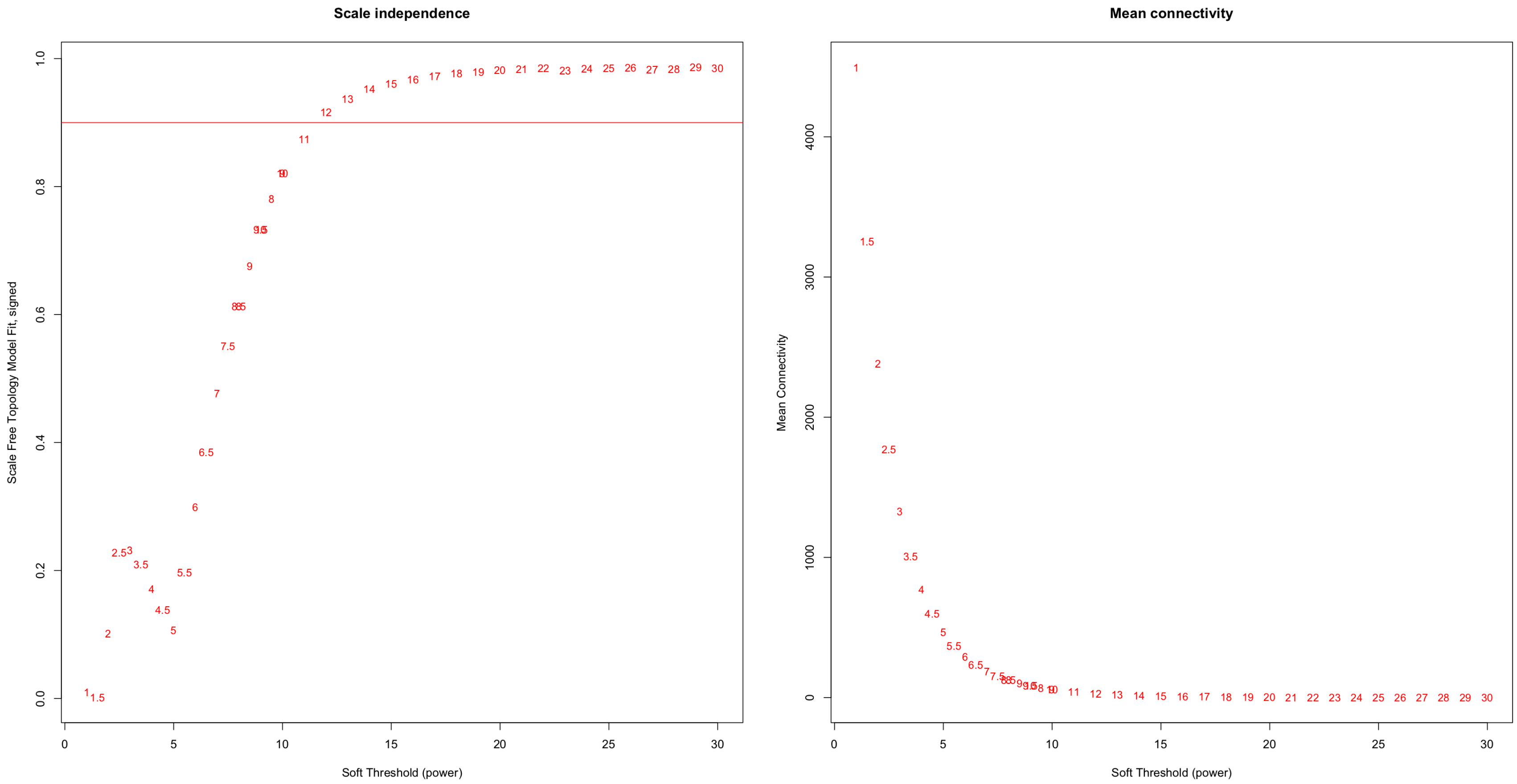


**Figure S2**. WGCNA scale independence and mean connectivity plots used for selection of soft threshold = 12. The red line represents $R^2 = 0.80$, or the point at which the network reaches scale-free topology.

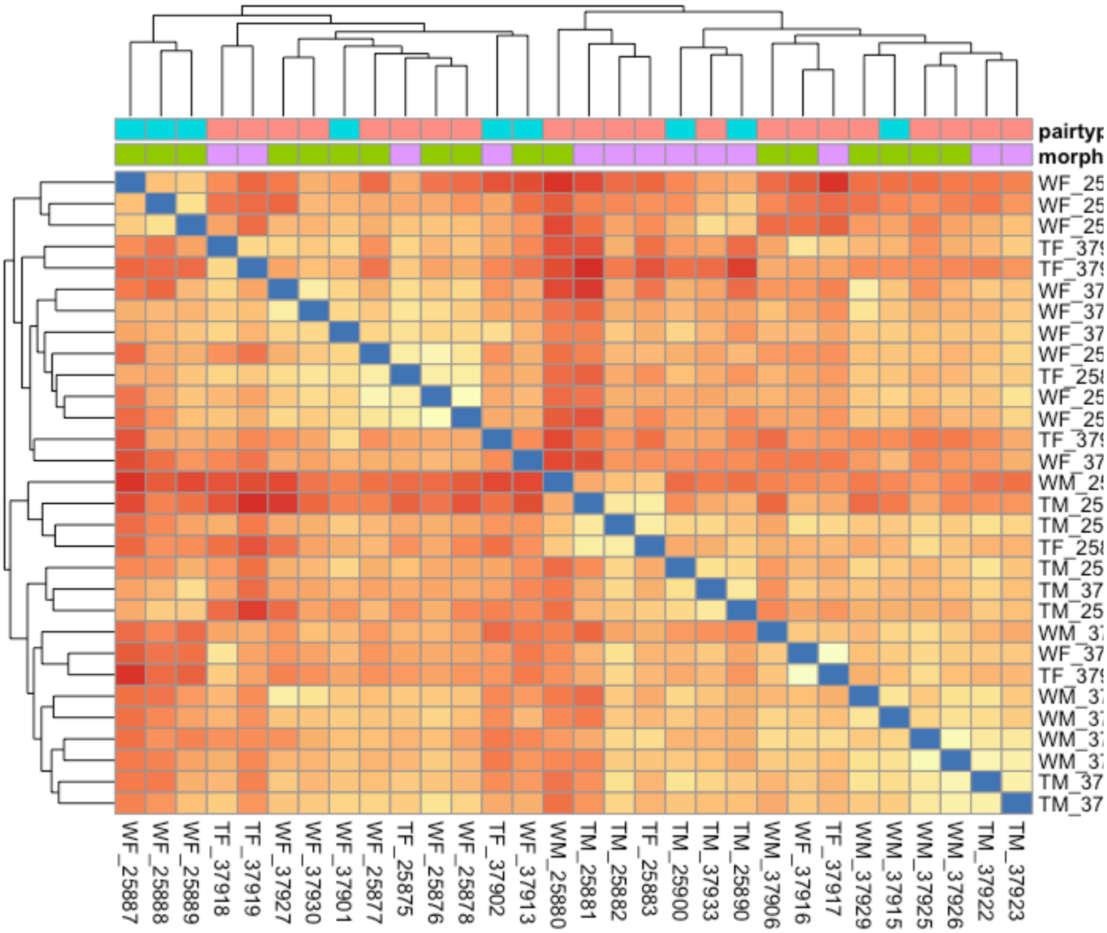


**Figure S3**. Hierarchical clustering heatmap showing relationships between samples, colored by morph and nest pairtype, based on all 8,982 genes used in differential expression testing.

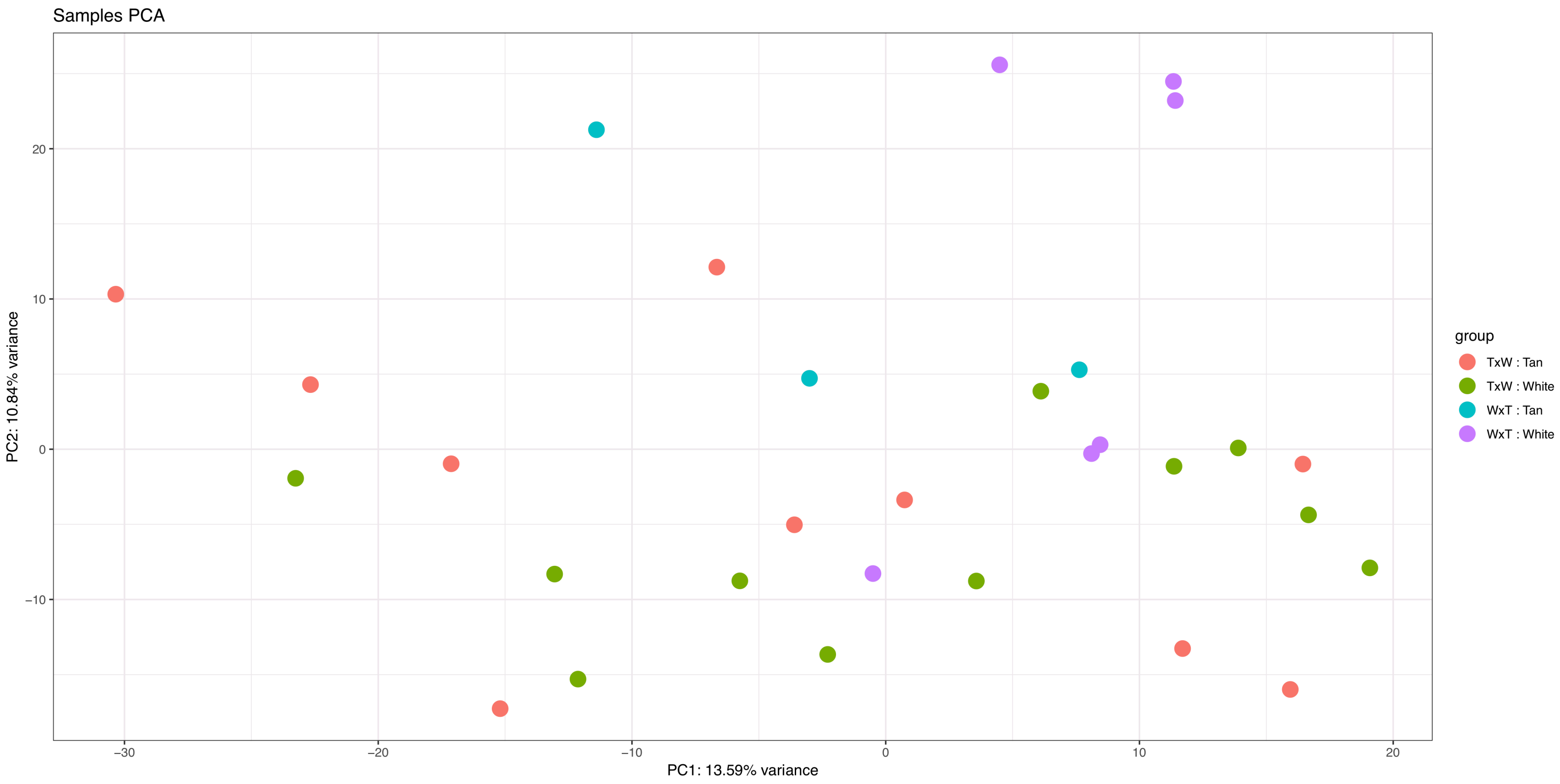


**Figure S4**. Principal components analysis (PCA) of all samples used in this study.

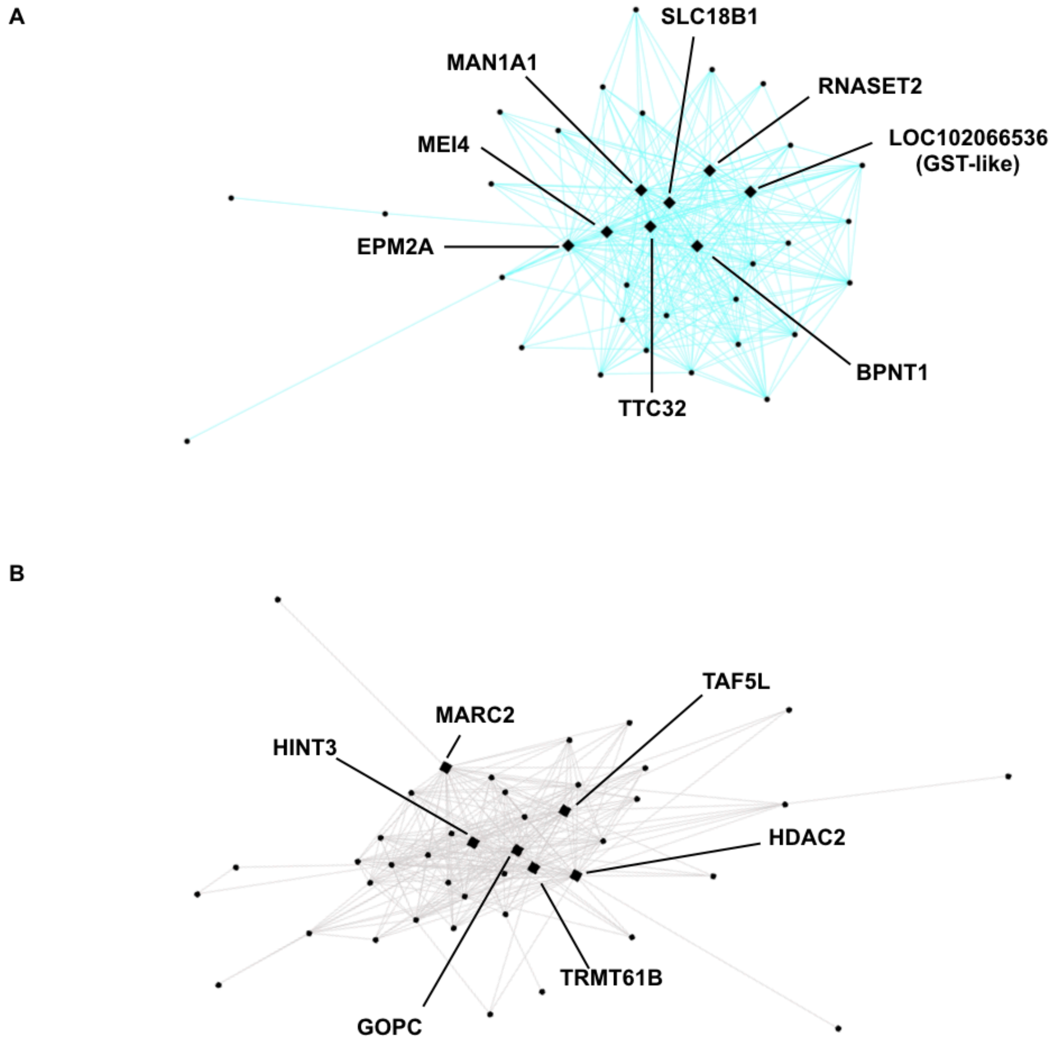


**Figure S5**. Networks of (A) light cyan and (B) ivory modules correlated with morph a driven by genes in ZAL2$^{m}$ inversion. Light cyan contains genes elevate and ivory contains genes suppressed in W morph nestlings. Each circle represents a gene and diamonds represent hub genes described in Table 1.

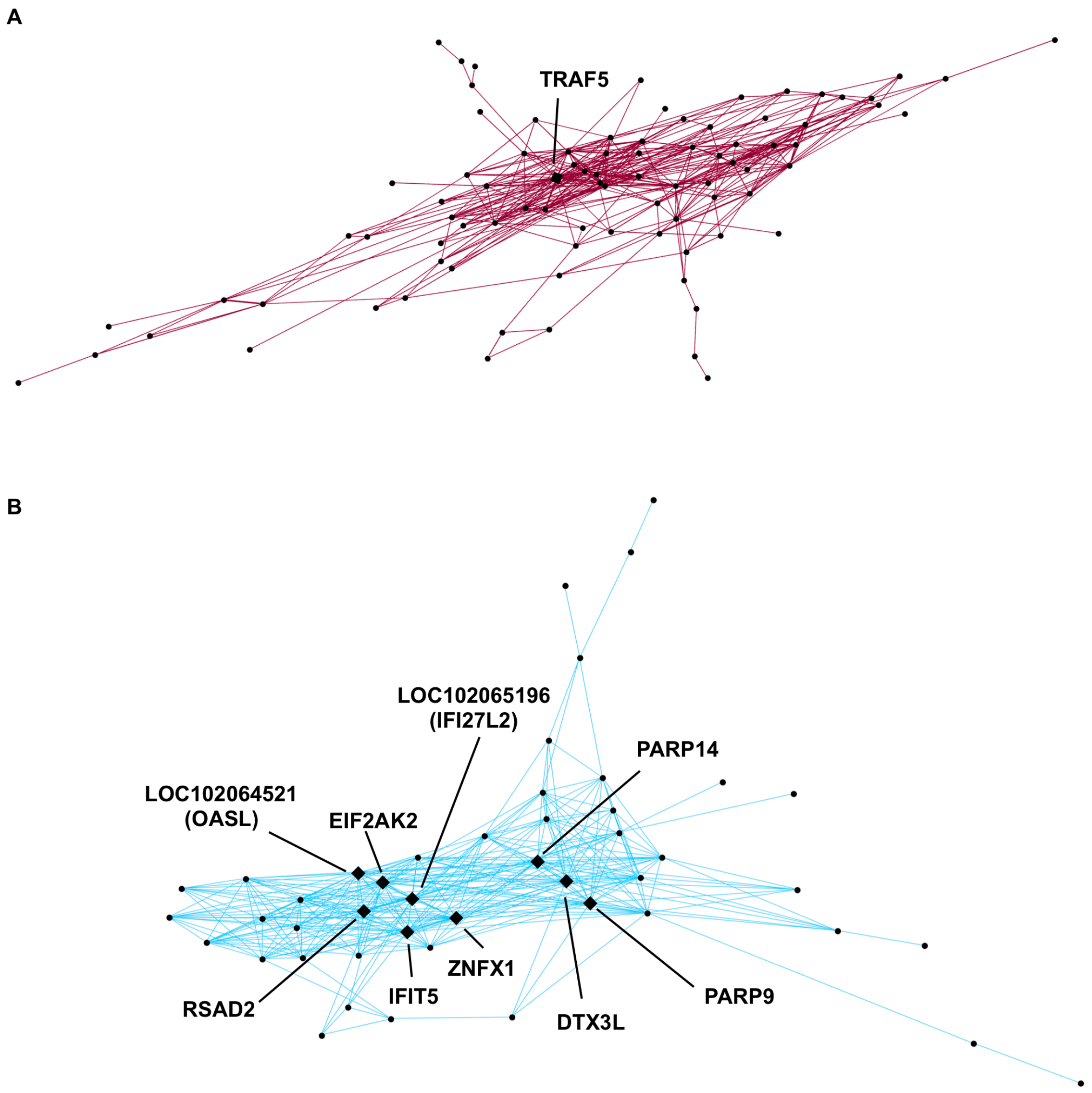


**Figure S6**. Networks of (A) dark red and (B) sky blue modules correlated with morph and driven by the increased expression of innate immunity hub genes. Each circle represents a gene and diamonds represent hub genes described in Table 1.

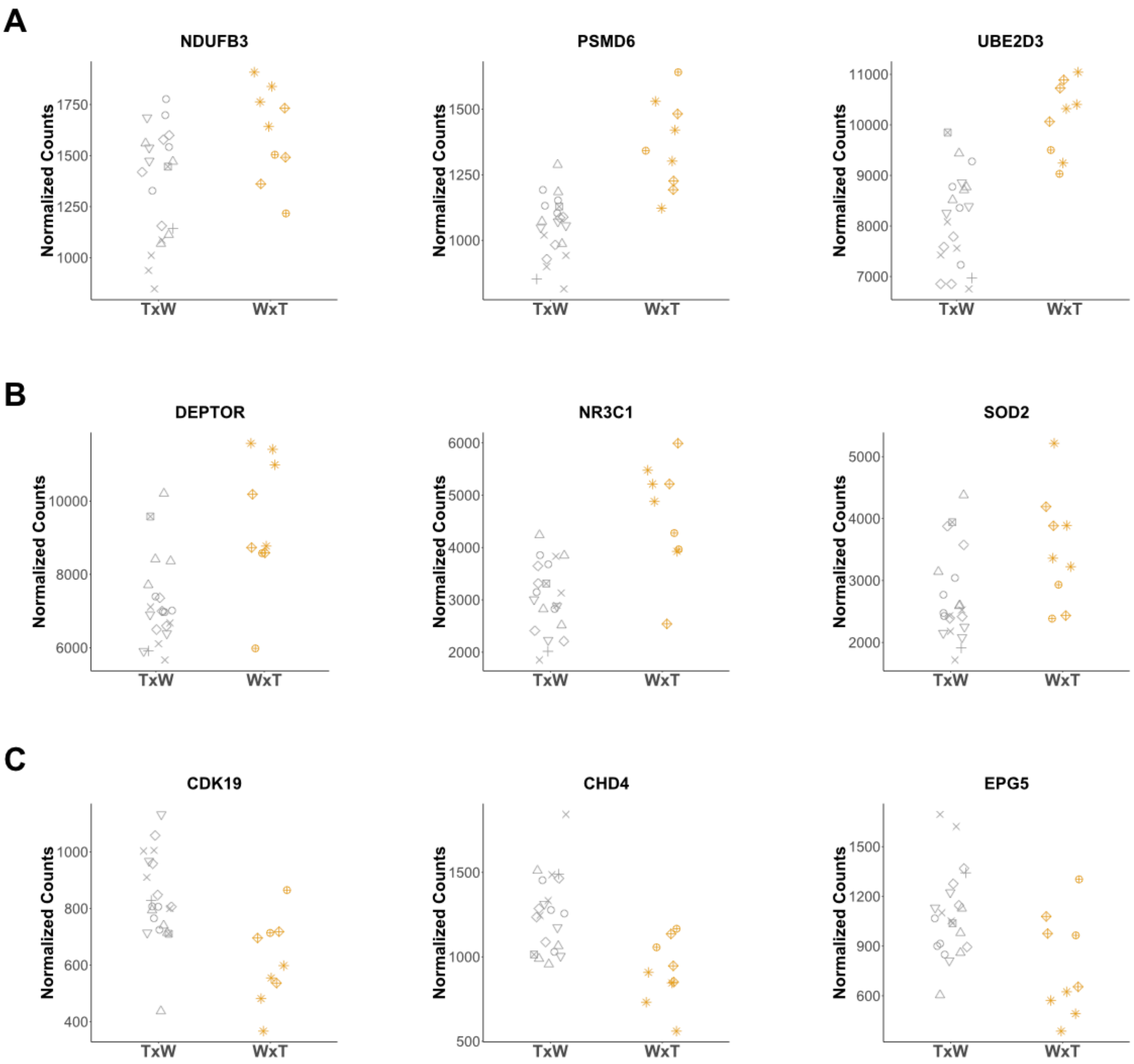


**Figure S7**. Normalized gene expression plots of candidate genes and/or hub genes found in (A) the blue module, (B) the beige module, and (C) the light green module. TxW represents samples from nests sired by a T male and a W female. WxT represent samples from nests sired by a W male and a T female. Plots are identical to those found in Figures 2-4, except the shape of points represents nest ID.
